# Amino acid supplementation confers protection to red blood cells prior to Plasmodium falciparum bystander stress

**DOI:** 10.1101/2023.05.16.540951

**Authors:** Heather Colvin Binns, Elmira Alipour, Dinah S. Nahid, John F. Whitesides, Anderson O’Brien Cox, Cristina M. Furdui, Glen S. Marrs, Daniel B. Kim-Shapiro, Regina Joice Cordy

## Abstract

Malaria is a highly oxidative parasitic disease in which anemia is the most common clinical symptom. A major contributor to malarial anemia pathogenesis is the destruction of bystander, uninfected red blood cells. Metabolic fluctuations are known to occur in the plasma of individuals with acute malaria, emphasizing the role of metabolic changes in disease progression and severity. Here, we report that conditioned media from *Plasmodium falciparum* culture induces oxidative stress in healthy uninfected RBCs. Additionally, we show the benefit of amino acid pre-exposure for RBCs and how this pre-treatment intrinsically prepares RBCs to mitigate oxidative stress.

**Key points:**

- Intracellular ROS is acquired in red blood cells incubated with *Plasmodium falciparum* conditioned media
- Glutamine, cysteine, and glycine amino acid supplementation increased glutathione biosynthesis and reduced ROS levels in stressed RBCs

## Introduction

Anemia is the most common clinical consequence of human malaria, a parasitic disease with nearly 250 million cases annually^1^. Pathogenesis of malarial anemia is multifaceted consisting of loss of both infected and uninfected red blood cells (RBCs) as well as dysregulation of the production of new RBCs into circulation^2^. In falciparum malaria, uninfected RBCs are lost at a much greater rate than infected RBCs. This bystander effect occurs prior to adaptive immune system activation^3^, contributing greatly to the development of malarial anemia. The mechanisms of malarial bystander effect on uninfected RBCs are unclear although loss in membrane deformability contributes to the removal of bystander RBCs from circulation^4, 5^. Bystander RBCs have also been shown to undergo cell surface changes promoting erythrophagocytosis through both complement-mediated activation^6^ and phosphatidylserine antibody-mediated removal^7^. Culture medium from *P. falciparum*-infected RBCs was shown to impact biological function of nucleated erythroid cells^8^. In addition, parasite-derived mitochondrial DNA within the media of *P. falciparum* parasite culture was found to elicit toll-like receptor 9 (TLR9) binding, thereby altering the membranes of healthy RBCs^9^. Interrupted glycolysis increases RBC susceptibility to senescence and oxidative damage and further highlights the importance of the exogenous metabolic environment for RBCs^10^. From this collective body of work, it is clear there are multiple mechanisms contributing to the bystander effect in malaria.

While mature RBCs have limited metabolic activity due to a lack of membrane-bound organelles, these cells have multiple active antioxidant components to counter oxidative stress in their environment. These defenses include reduced glutathione (GSH), catalase, peroxiredoxins, and glutathione peroxidase^11, 12^. Oxidative stress plays a major role in many anemia-inducing conditions, such as malaria and sickle cell disease^13, 14^. In malaria, RBCs from patients have reduced levels of intracellular catalase^15^, indicating these cells are deficient in their ability of combatting oxidative stress and rendering avenues of antioxidant therapy as a viable treatment to lessen disease severity. Additionally, significant host metabolic alterations occur during malaria. This includes markedly reduced levels of plasma free amino acids such as glutamine and arginine^16–23^, suggesting amino acid supplementation could perhaps provide therapeutic benefit in malaria.

Oral treatment with L-glutamine (Gln) is a recently approved therapy for sickle cell disease^24^. While the exact cellular mechanism remains unclear, a potential role for Gln is to improve the nicotinamide adenine dinucleotide phosphate (NADPH) stores in sickled RBCs^25^, lessening the oxidative stress. Gln is also implicated in malarial anemia where low plasma Gln levels were found to be associated with severe pediatric malarial anemia^26^. Here, we explore the oxidative impact of the *P. falciparum* culture environment on uninfected RBCs as a proxy for *in vivo* malaria bystander effect. Furthermore, we investigate the role of exogenous amino acid supplementation on bystander RBCs and show that RBCs pretreated with precursor antioxidant amino acids glutamine, cysteine and glycine have increased intracellular glutathione synthesis and thus confer protection from oxidative stress.

## Methods

### Blood washing and perturbations

Heparinized venous blood, Type O+, was collected from healthy adult participants with informed consent and used with approval from Wake Forest University Institutional Review Board (Study ID: IRB00024199) or commercially purchased from BioIVT Inc. Briefly, RBCs were washed three times in excess 1X PBS pH 7.4 (Gibco™) and centrifuged at 900 x g to remove plasma and buffy coat layers with each wash. Washed RBCs were stored in 1X PBS supplemented with 5 mM D-glucose (PBS+Gluc) and used same day. All RBC incubation experiments were performed at 1% hematocrit (hct) while rocking at 37°C. Overnight 24-hour incubations were performed for amino acid supplementation and *P. falciparum* conditioned medium (*Pf*CM) stress. For amino acid supplementation experiments, indicated concentrations of amino acids were thoroughly dissolved in PBS+Gluc and incubated overnight. Cells were then washed twice in PBS to remove amino acids prior to stress incubations. All hydrogen peroxide perturbations were performed in the presence of 1 mM sodium azide to block intracellular catalase activity and incubated for 15-minutes while rocking at 37°C, as described previously^27–29^. *Pf*CM perturbations were also performed in the presence of 1 mM sodium azide to block intracellular catalase activity.

### Plasmodium falciparum culture and conditioned media

*Plasmodium falciparum* of the 3D7 laboratory strain (MRA-102; BEI Resources) was utilized in the generation of *Pf*CM. The parasites were cultured in a plugged 75cm^2^ flask at a total volume of 25 mL with RBCs combined from two O+ donors (BioIVT Inc.) at 2% hct and standard culture media consisting of RPMI 1640 (Gibco™) supplemented with sodium bicarbonate, HEPES buffer, hypoxanthine, gentamicin, and 0.5% (wt/vol) Albumax II^30^. A filtered gas line connected to a gas tank containing 1% O_2_, 5% CO_2_, and 94% N_2_ gas mixture was placed inside the flask for about 45 seconds to induce the deoxygenated environment needed for parasite growth. This gas mixture was administered every 48 hours for maintaining the optimal oxygen environment in the flask during the maintenance period. In preparation for *Pf*CM media generation, parasites were synchronized using a 5% Sorbitol solution at least two cycle days prior to culture and subsequent *Pf*CM collection. Synchronized parasites were seeded at 0.4% parasitemia and followed for 72 hours; at 48 hours the culture was re-gassed. The culture was then centrifuged at 600 x g for 15 minutes, followed by 1600 x g and 3600 x g, as described previously^31^. Collected *Pf*CM was filtered with a 0.45 μm filter to further remove any remaining cellular debris. Filtered *Pf*CM was kept at 4°C for downstream use on the same day of generation.

### Intracellular ROS detection

Intracellular ROS was measured using a Cellular ROS Assay Kit (ab113851; Abcam). Briefly, washed RBCs were incubated with a 2 µM working concentration of 2’,7’ – dichlorofluorescin diacetate (DCFDA) for 30 minutes. RBCs were then washed twice with 1X PBS and used for either flow cytometry analysis or live cell imaging.

### Flow cytometry

RBCs were analyzed by a BD LSRFortessa™ X-20 Flow Cytometer (Becton Dickinson). RBCs were passed at a flow rate of about 5,000 cells per second. A 60 milliwatt 488 nm laser at 550 volts was used for excitation and the green (505 longpass and 530/30 bandpass), fluorescein isothiocyanate (FITC) emission filter was used for detection. For data analysis, mean fluorescent intensity was measured for 100,000 events, gated on doublet discrimination (FSC-H vs FSC-A), and analyzed using FCS Express 7 Research software (De Novo).

### Live cell imaging

Experiments were carried out using a Leica Thunder Live Cell Imaging System and LasX acquisition software. Images were acquired as 16-bit data with a Leica K8 sCMOS camera (2048 X 2048 pixels), using a 63X Plan Apo oil immersion lens (1.4NA). RBC cells were plated onto 35 mm optical coverglass dishes (Ibidi) immediately before imaging. For time lapse experiments, images were acquired every 15 seconds. All excitation/acquisition parameters were carefully held constant across imaging experiments including light-emitting diode excitation level and camera exposure time. All analyses were performed using Leica instantaneous computation clearing intensity (ICC) values. For analysis, stationery cells were chosen using Differential Interference Contrast images and isolated using region of interest selection using FIJI^32^. Fluorescence in regions of interest was quantified from ICC adjusted images at each timeframe image. Data was normalized to average fluorescence of first three timepoints, prior to treatment. For visualization purposes, images were adjusted to maximize for brightness contrast using FIJI^32^.

### Scanning electron microscopy

Perturbed RBCs were washed and fixed in an osmotic-controlled glutaraldehyde solution, as previously reported^33^. Briefly, glutaraldehyde-fixed RBCs were washed and resuspended in distilled H_2_O at 0.5% hct before air drying overnight on 12mm round coverslips at 60°C. Images were collected with Everhart-Thornley secondary electron detection using either an AMRAY 1810 or a Phenom XL scanning electron microscope at 10Kv accelerating voltage. Typical magnifications employed were 2000X to allow for high resolution of RBCs while maintaining a reasonable field size. Samples were prepared for imaging by first dehydrating on carbon tab aluminum stubs, then gold sputter coating under argon gas conditions to a thickness of about 7-10nm. Echinocyte morphology stages were defined as previously described^34^ and quantified per 100 cells per sample slide.

### Osmotic gradient ektacytometry

A Technicon osmotic gradient ektacytometer (Technicon Instrument Corp.) facilitated the deformability measurements of the RBCs. Thirty-one (31) g/L of polyvinylpyrrolidone (PVP) polymer (Sigma, 437190) mixed with 0.9 g/L of sodium phosphate dibasic anhydrous Na_2_HPO_4_ (Fisher, 7558-79-4), 0.24 g/L of sodium phosphate monobasic NaH_2_PO_4_ (Fisher Biotech, 10049-21-5), and 0.544 g/L of sodium chloride NaCl (Sigma, S7653) was prepared in Milli-Q ultrapure water with the pH of 7.4. The “low” solution (40 mOsm) was used to make the “high” (750 mOsm) and “sample” solutions, by dissolving 11.25 g of NaCl in 500 mL of the low solution, and 1.9782 g of NaCl in 250 mL of the low solution (290 mOsm), respectively. After calibration using known osmolality mixtures and finding the required parameters as well as laser alignment of the ektacytometer, 150 μL of the blood samples (40% hematocrit) was diluted into 4 mL of sample solution. The population of RBCs suspended in high viscous media was directed into the gap between two coaxial cylinders. The outer cylinder stayed motionless while the inner cylinder rotated with distinct angular velocity to apply the defined shear stress of 159 dynes/cm^2^ (∼16 pascals) at a controlled temperature. During operation, the focused laser beam passed through the suspension and generated elliptical diffraction patterns of the flowing RBCs projected on a detector. LabVIEW software recorded the diffraction patterns and the corresponding osmoscans and quantitative statistics. Data was fit using Origin 2016 software and the averages of three separate scans from each blood sample were analyzed. GraphPad Prism was used for the graphing of the recorded result.

### Metabolite extraction

Following perturbation experiments, 1×10^7^ RBCs were pelleted and the supernatant was carefully removed. Pelleted cells were resuspended in 1X PBS with 100 mM N-ethylmaleimide (NEM) (Cat#128-53-0, Millipore Sigma) for 15 minutes at room temperature^27^. Following NEM incubation, cells were washed twice with 1X PBS. After thorough removal of NEM from supernatant, PBS was added with a final cell concentration of 20% hct and cell concentration was recorded using a hemocytometer. Methanol was added (4:1) followed by vortexing and storing cells on ice for 30 minutes. Sample tubes were then centrifuged at 18,000 x g and supernatant was carefully removed and stored in −80°C for mass spectrometry analysis.

### Targeted Mass Spectrometry

Targeted LC-MS/MS analysis was performed at the Proteomics and Metabolomics Shared Resource (Wake Forest University School of Medicine, Winston Salem, NC). Briefly, extracted samples were dehydrated and reconstituted in H_2_O followed by mass spectrometry (Sciex 7500 MS) analysis^35^ for relative quantification of reduced glutathione (GSH alkylated by NEM) and oxidized glutathione (GSSG) metabolites, without any further derivatization^36^. Separation was performed on a Thermo Scientific Hypersil GOLD aQ reverse phase column (2.1 x 150 mm, 3µm) with a gradient mobile phase system consisting of an aqueous phase of 0.1% formic acid (A) and an organic phase of acetonitrile (B) at a flow rate of 0.5 mL/min (0 - 0.5 min, 0.5-5% B; 0.5 - 6.5 min, 5-98% B; 6.5 min-9 min, 98% B). The mass spectrometer used the following source parameters: Ion source gas 1: 35 psi, Ion source gas 2: 70 psi, Curtain gas: 40 psi, CAD gas: 9 psi, Source temperature: 250°C, Spray voltage: 5500 V. Transition masses for targeted analysis were 433.00 > 304.00 m/z (NEM-labeled GSH), and 613.20 > 355.25, 613.20 > 484.20 and 613.20 > 231.05 m/z for GSSG. Relative peak intensity values were normalized to 10^6^ cells per sample. Total glutathione levels were determined by adding intensities of GSH-NEM and GSSG with correction for ionization efficiency.

## Results

### Exogenous Gln alone is not sufficient but exogenous amino acid cocktail reduces oxidative stress acquired from H_2_O_2_

To assess the impact of amino acid supplementation on RBCs in highly oxidative diseases, we pre-exposed RBCs to amino acids recapitulating increased plasma concentrations that might result from oral supplementation. Following supplementation, cells were washed twice to remove amino acids from the exogenous environment and then exposed to 50 µM hydrogen peroxide (H_2_O_2_) and 1 mM sodium azide (NaN_3_) to induce oxidative stress^27^. We first sought to explore the impact of exogenous Gln supplementation on RBCs when exposed to oxidative stress, given its relevance in both malaria and sickle cell anemia. To do so, we measured intracellular reactive oxygen species (ROS) via 2’,7’ –dichlorofluorescin diacetate (DCFDA). Interestingly, though we found a significant difference in intracellular ROS levels upon H_2_O_2_ treatment (fold change of 1.0 for 0 µM H_2_O_2_ vs 4.1 for 50 µM H_2_O_2_, p = 0.036), we found no difference between H_2_O_2_-stressed RBCs pre-exposed to control media or media supplemented with Gln overnight (**Figure 1A**).

**Figure 1:**
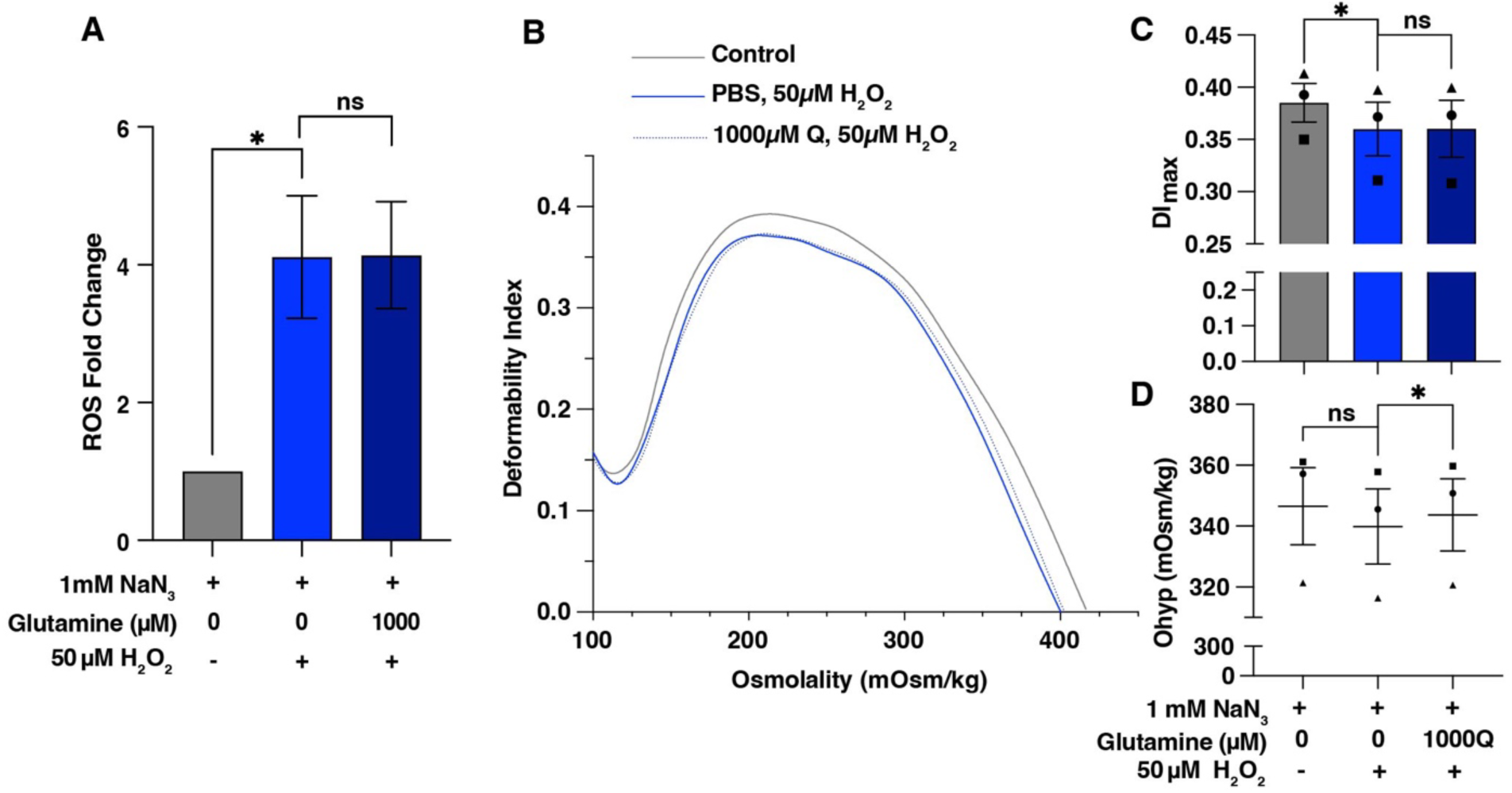
Glutamine pre-treatment benefitted oxidatively stressed RBCs through osmotic protection. (**A**) Fold change of intracellular ROS detected by DCFDA staining and flow cytometry in RBCs. (**B**) Representative ektacytometry curve as cells pass through osmotic gradient at a constant shear stress of 16 Pa. (**C**) Maximum deformability index (DI_max_) values and (**D**) RBC hydration graphed as O_hyper_.; n=3. Mean ± standard error of the mean denoted by error bars. One-way paired t-test; ns = not significant, \**p < 0.05*

As H_2_O_2_ is known to impair RBC function through reduced RBC membrane deformability and cellular dehydration^28, 29, 37^, we next measured biophysical properties of the RBCs. To determine the osmotic effect of Gln supplementation, we utilized ektacytometry which measures the diffraction pattern of a laser hitting a population of RBCs at a constant shear stress while osmolality increases.^38^ Here, we saw an expected decrease in membrane deformability, as measured by DI_max_, in response to H_2_O_2_ (0.385 DI_max_ for 0 µM H_2_O_2_ vs 0.360 DI_max_ 50 µM H_2_O_2_, p = 0.035). However, RBCs supplemented with Gln had a similar loss in deformability when stressed with H_2_O_2_, suggesting no impact of Gln on that parameter (**Figure 1B and 1C**).

In terms of cell hydration, we observed slight dehydration in RBCs stressed by H_2_O_2_ that bordered on significance (347 mOsm/kg for 0 µM H_2_O_2_ vs 340 mOsm/kg for 50 µM H_2_O_2_, p = 0.061), but more interestingly, we found that Gln supplementation significantly improved hydration status compared to RBCs not supplemented with Gln (340 mOsm/kg for 0 µM Gln vs 344 mOsm/kg 1000 µM Gln, p = 0.031) (**Figure 1D**). These data show that Gln supplementation prior to oxidative stress was osmotically advantageous for RBCs hydration but did not improve RBCs membrane deformability nor reduce intracellular ROS generation in RBCs.

Although Gln supplementation alone did not seem to protect RBCs from oxidative stress, Gln is a precursor for glutamate which is required, in addition to cysteine and glycine, for *de novo* GSH synthesis in RBCs to combat high levels of oxidative stress^39, 40^. Therefore, to be sure all components of this pathway were available exogenously, we pretreated RBCs with media supplemented with Gln (Q), cysteine (C), and glycine (G), hereby referred to as QCG, to determine if these exogenous amino acids could equip RBCs to mitigate oxidative stress. Indeed, we found that RBCs supplemented with 1000 µM QCG amino acids incurred significantly less intracellular ROS when stressed with H_2_O_2_ (mean fold change of 5.0 without QCG vs 4.0 with 1000 µM QCG pre-treatment, p = 0.009) (**Figure 2A**). This oxidative protection was dose-dependent with significant protection conferred from 1000 µM QCG supplementation. We also found RBCs pre-exposed to QCG had a dose-dependent reduction in loss of membrane deformability once stressed with H_2_O_2_ (**Figures 2B-C**), although the 1000 µM QCG concentration of pretreatment did not reach significance (p = 0.068). Similar to Gln supplementation, QCG pretreatment resulted in significantly higher hydration status in RBCs (340 mOsm/kg without QCG vs 350 mOsm/kg with QCG, p=0.021), as measured by O_hyp_ (**Figures 2B and 2D**). Thus, we found that RBCs pretreated with QCG amino acids were partially protected from H_2_O_2_ induced oxidative stress.

**Figure 2:**
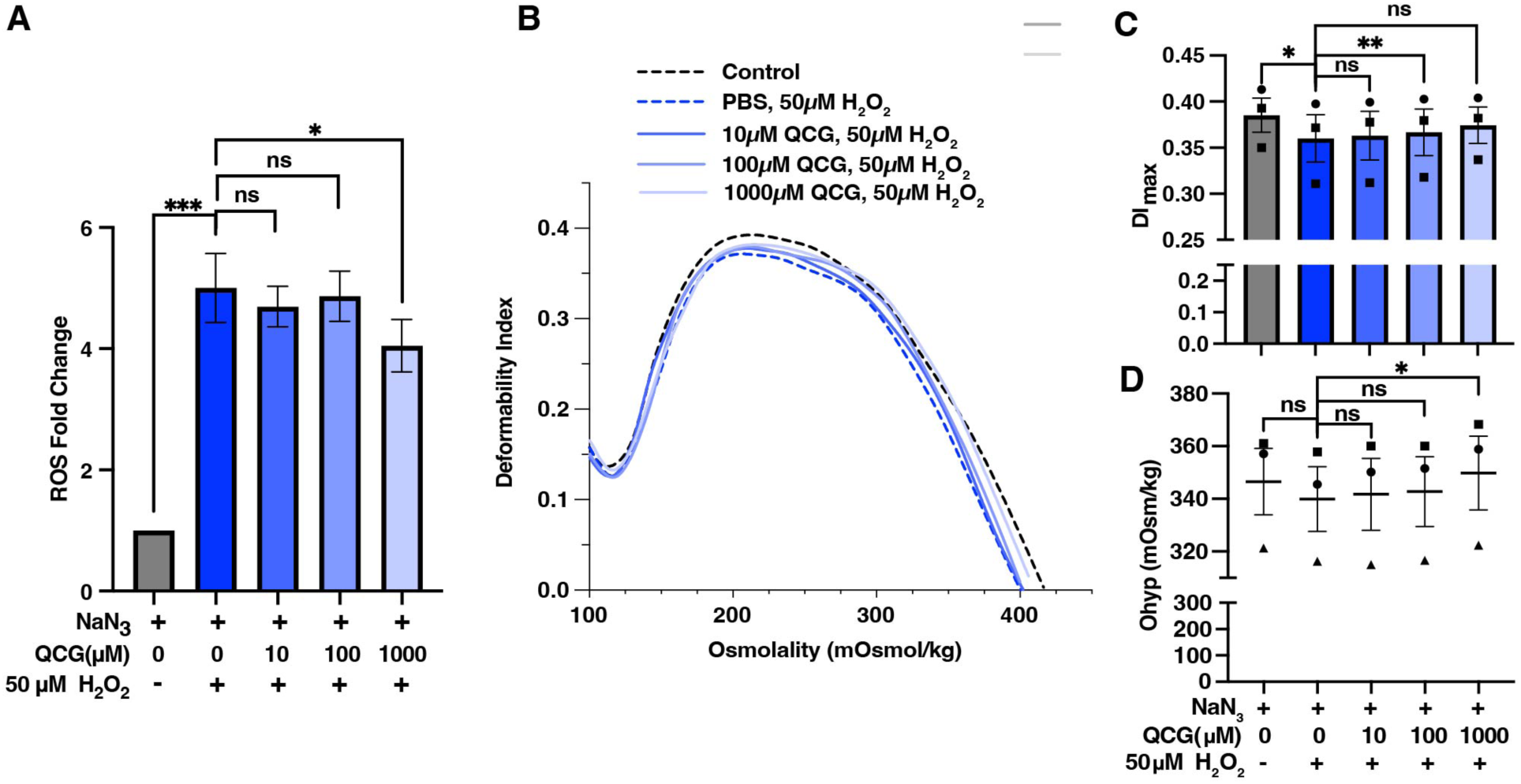
QCG preincubation reduces oxidative impact of H_2_O_2_ induced stress. (**A**) Fold change of intracellular ROS detected by DCFDA staining and flow cytometry in RBCs; n=6 (**B**) Representative ektacytometry curve as cells pass through osmotic gradient at a constant shear stress of 16 Pa. (**C**) Maximum deformability and (**D**) hydration level of RBCs from ektacytometry curves; n=3. Mean ± standard error of the mean denoted by error bars. One-way paired t-test; ns = not significant, \**p < 0.05, ** p < 0.005, *** p<0.0005*.

### *P. falciparum*-conditioned medium induces stress in human RBCs

To determine the oxidative impact of *P. falciparum* conditioned medium (*Pf*CM) on uninfected RBCs, we exposed human RBCs from healthy participants to *Pf*CM in the presence of 1mM NaN_3_ and measured RBC echinocytosis, a morphological change associated with oxidative stress^41^. Utilizing scanning electron microscopy (SEM), we scored RBCs on the severity of echinocyte morphology **(Supplemental Figure 1A)** as previously reported^34, 42, 43^. We observed that RBCs exposed to *Pf*CM had significantly higher morphology scores (mean score of 116 for control vs. 138 for *Pf*CM, p = 0.0002) (**Figure 3A**) and a higher percentage of echinocytes (mean of 10.5% for control vs. 24.9% for *Pf*CM, p = 0.002) (**Figure 3B**), as compared to RBCs in control RPMI media. Although overall RBC morphology stages showed slight variations between participants, more severe echinocyte stages were present in RBCs from all participants exposed to *Pf*CM **(Supplemental Figure 1B)**. Next, we aimed to determine whether redox state was affected in mature RBCs alongside the change in morphology. RBCs incubated with *Pf*CM had a significant increase in intracellular ROS as detected by DCFDA, compared to RBCs from the same donor exposed to control media (fold change of 1.0 for control vs. 1.8 for *Pf*CM, p = 0.011) (**Figure 3C-D**). Thus, *Pf*CM induces oxidative stress in uninfected RBCs, suggesting a role for oxidative stress in malaria bystander effect.

**Figure 3:**
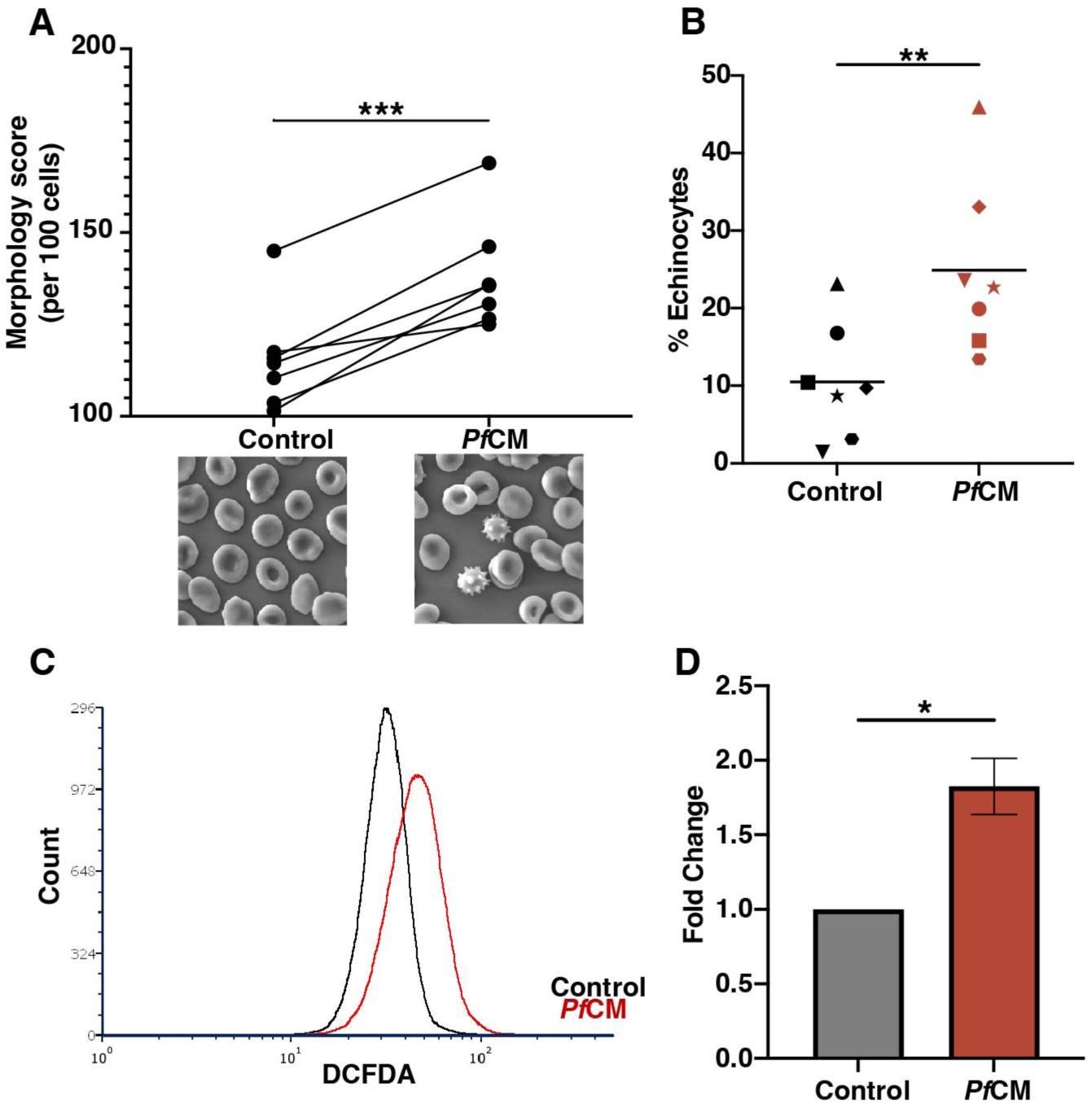
*Pf*CM increases RBC echinocytosis and intracellular ROS. RBCs incubated in control media or *Pf*CM overnight were imaged with SEM and assessed based on (**A**) morphology scores and (**B**) percentage echinocytes (n=7). RBCs incubated in control media or *Pf*CM overnight were washed after incubations and stained with DCFDA to quantify intracellular ROS with flow cytometry. Representative flow cytometry histogram shown in (**C**) and bar graph of fluorescence fold change compared to control in (**D**); n=4. Mean ± standard error of the mean denoted by error bars. One-way paired t-test; *p < 0.05, ** p < 0.005, *** p<0.0005.

### QCG supplementation reduces *Pf*CM induced oxidative stress in RBCs

We next aimed to determine whether QCG supplementation also conferred protection to RBCs exposed to *Pf*CM. Indeed, RBCs pretreated with QCG amino acids were found to have a significant reduction in intracellular ROS following *Pf*CM incubation (mean fold change of 1.72 for 0 µM QCG vs. 1.15 for 1000 µM QCG, p = 0.027) (**Figure 4A**). Morphologically, we observed RBCs pretreated with QCG had overall improved morphology scores after *Pf*CM stress (mean score of 130 for 0 µM QCG vs. 122 for 1000 µM QCG, p = 0.040) (**Figure 4B**). Therefore, we found that QCG supplementation confers protection to *Pf*CM induced oxidative stress in RBCs.

**Figure 4:**
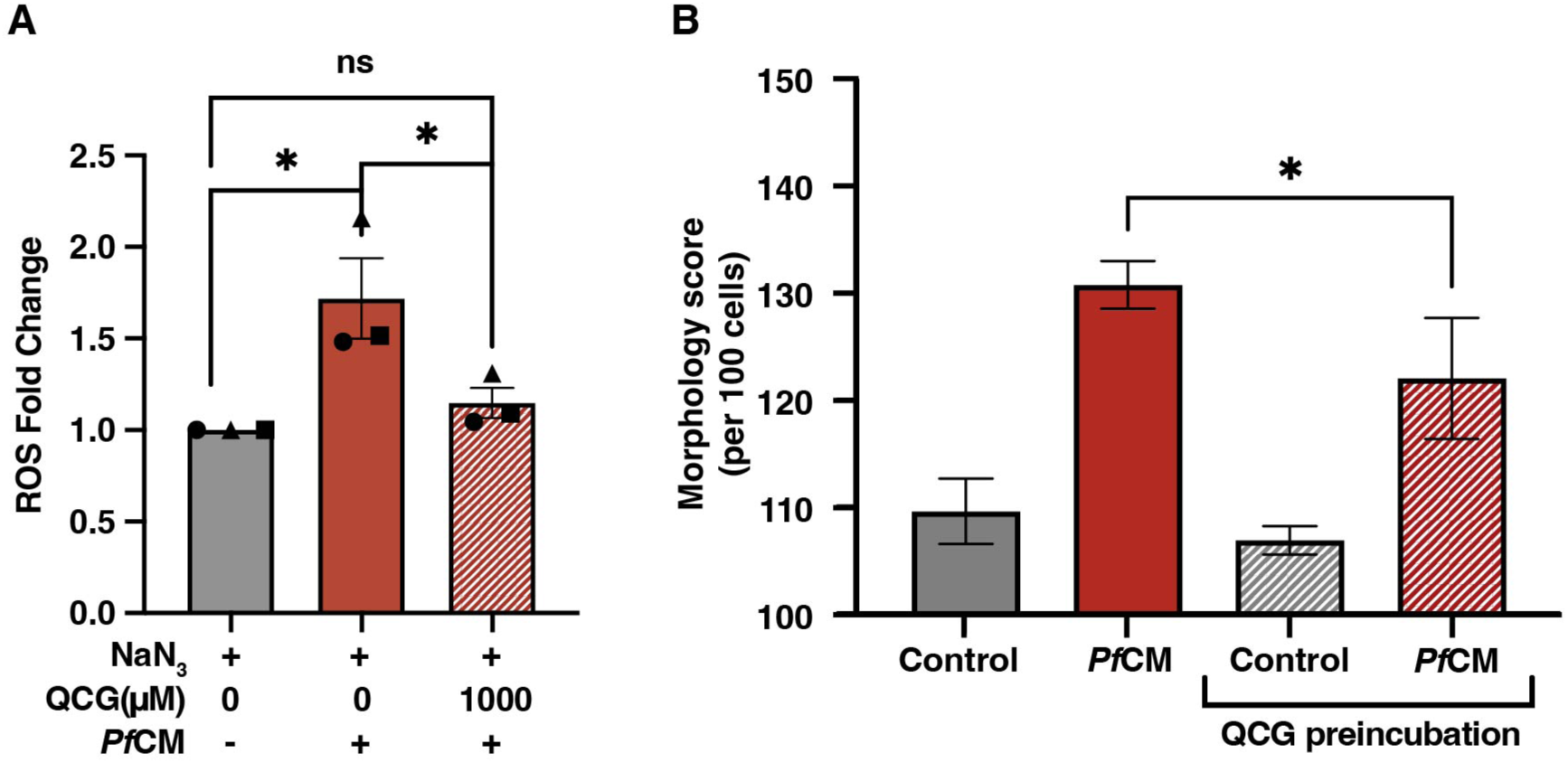
QCG supplementation protects RBCs from *Pf*CM induced oxidative stress. (**A**) Fold change of intracellular ROS detected by DCFDA staining and flow cytometry in RBCs, n=3. (**B**) RBC morphology score assessed from scanning electron microscopy images, n=5. Mean ± standard error of the mean denoted by error bars. One-way paired t-test; ns = not significant, \**p < 0.05*

### QCG supplementation induces intracellular RBC glutathione synthesis

We hypothesized that QCG protection occurs via intracellular GSH metabolic pathways given that each amino acid is a known precursor for *de novo* GSH synthesis (**Figure 5A**). To determine the metabolic effect of QCG supplementation and test this hypothesis, we performed targeted metabolomics (LC-MS/MS) on RBCs preincubated with QCG or PBS (control) and then stressed with either *Pf*CM or H_2_O_2_ (50µM). We found increased levels of total glutathione in QCG supplemented RBCs (**Figures 5B-D**, **Supplementary Figures 2A-C**). This activity was evident from QCG supplementation, with or without induced oxidative stress. We also observed an increased level of oxidized glutathione (GSSG) in QCG supplemented RBCs after exposure to either *Pf*CM (1525 mean peak area intensity for 0 µM QCG vs 6725 for 1000 µM QCG, p = 0.009) (**Figure 5C**) or H_2_O_2_ (111,650 mean peak area intensity for 0 µM QCG vs 211,245 for 1000 µM QCG, p = 0.103) (**Supplemental Figure 2B)** compared to RBCs without QCG supplementation or exposure to oxidative stress. Thus, the results indicate that QCG supplemented RBCs have increased intracellular glutathione biosynthesis.

**Figure 5:**
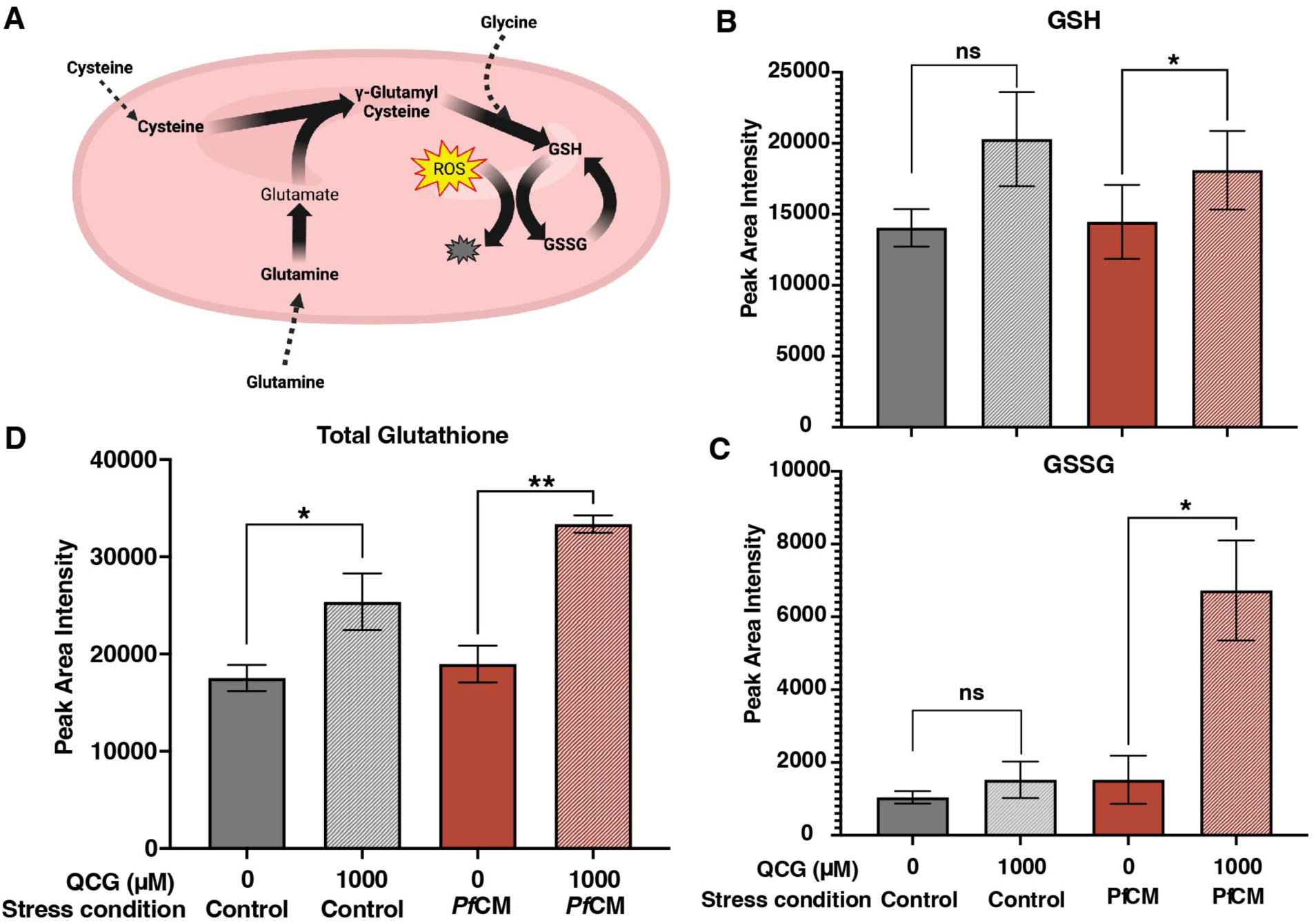
Glutathione metabolic changes within RBCs supplemented with QCG followed by *Pf*CM stress. (**A**) Schematic of metabolic processes known to occur within RBCs from QCG amino acids. Peak area intensity of (**B**) reduced glutathione (GSH), (**C**) oxidized glutathione (GSSG), and (**D**) total glutathione as measured by LC-MS/MS, n=3. One-way paired t-test; ns = not significant, \**p < 0.05, ** p < 0.005*.

### Supplementation with QCG promotes RBC intrinsic antioxidant properties

Given the aforementioned findings, it remained unclear whether QCG-mediated protection was a generalized RBC response or if QCG protection occurred specifically under oxidative stress exposure. To determine this, we analyzed the kinetic response of QCG supplemented RBCs to oxidative stress. Using fluorescence imaging, we measured oxidation in RBCs containing H_2_DCF and exposed to 50µM H_2_O_2_ over 5 min. RBCs pre-exposed to QCG showed reduced levels of intracellular ROS within 2.5 min of H_2_O_2_-induced oxidative stress while RBCs pre-exposed to either PBS or Gln had higher detected ROS that developed quicker within the cells (**Figure 6A**). Cellular response to oxidative stress among all preincubation conditions appeared to be heterogenous across cells (**Figures 6B-D**). Interestingly, we observed punctate-like areas of increased fluorescence in each preincubation condition with or without H_2_O_2_-induced oxidative stress suggesting some internal organization of intracellular ROS. This phenomenon was observed in both H_2_O_2_ and *Pf*CM-stressed RBCs (**Figure 6E**). Together, these data indicate QCG preincubation intrinsically prepares RBCs to counter oxidative stress prior to any stress exposure rather than mounting an antioxidant response following oxidative stress.

**Figure 6:**
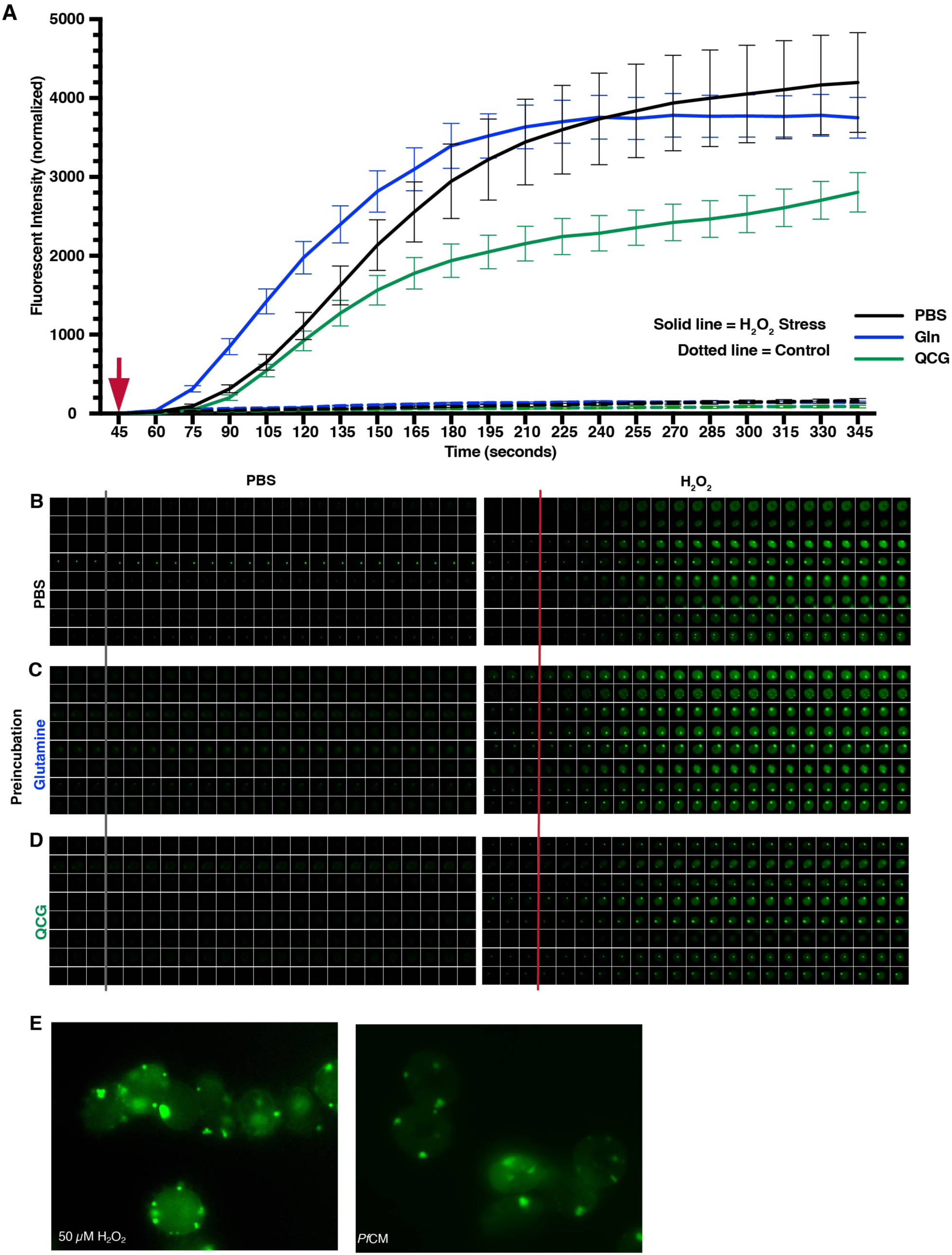
QCG supplementation rapidly protects RBCs from intracellular ROS formation during oxidative stress. RBCs pretreated with either PBS, Gln (Q), or QCG were labeled with DCFDA and placed in a glass bottom petri dish per sample condition and allowed to settle for 45 seconds before given a bolus (red arrow, **A**) of PBS (grey line, **B-D**) or 50 µM H_2_O_2_ (red line, **B-D**). Stationary cells (total 8 cells per condition) were quantified (**A**) and imaged (**B-D**) per condition at each 15 second time point for five minutes. Images were taken with Leica Thunder at 63X oil immersion (**A-D**). Representative images of DCFDA labeled RBCs stressed with 50 µM H_2_O_2_ (left) or *Pf*CM (right) (**E**). Images taken with Echo Revolve Fluorescent Microscope at 100X oil immersion (**E**).

## Discussion

In this study, we investigated the metabolic role of amino acid supplementation on oxidatively stressed RBCs with an overall goal of alleviating RBC oxidative burden. We first explored Gln supplementation on RBCs stressed by H_2_O_2_ and observed that while Gln supplementation alone was not sufficient to reduce oxidative stress levels within RBCs, Gln supplementation promoted RBC hydration (**Figure 1**). In contrast, we identified a significant decrease in intracellular ROS in RBCs that had been supplemented with QCG amino acids (**Figure 2**). These data suggest that observed ROS protection is conferred through RBC GSH biosynthesis pathway, as total glutathione was increased in RBCs supplemented with QCG (**Figure 5**) and this protection from intracellular ROS development was found to be rapid (2.5 min) upon induction of oxidative stress (**Figure 6**). Here, we also report for the first time to our knowledge that conditioned media from *P. falciparum* culture increases intracellular ROS in uninfected RBCs (**Figure 3**), and is a likely contributor to malaria bystander effect. Additionally, we found that QCG supplementation does indeed confer protection in the form of reduced intracellular ROS and improved echinocyte morphology to *Pf*CM stressed RBCs (**Figure 4**), highlighting a therapeutic prospect for malarial anemia.

Recently *Pf*CM was shown to increase oxidative stress in erythroid precursor cells^8^ and alter membrane structure and binding in uninfected mature RBCs^9^. However, it was unclear if *Pf*CM also perturbed the oxidative status of uninfected mature RBCs. We confirmed one mode of action that *Pf*CM has on mature RBCs is through oxidative stress as measured by the induction of intracellular ROS. In general, elevated ROS reduces RBC survivability *in vivo*^44^, suggesting this could be an additional contributor to pathogenesis of bystander effect in malaria.

Gln is implicated in both sickle cell anemia and malarial anemia, highlighting a possible role for metabolic intervention in anemic conditions. Lower plasma Gln levels are associated with pediatric malarial anemia^26^ while oral Gln supplementation is an approved treatment for sickle cell anemia, although cellular mechanisms of this therapy are still under investigation. Here, we show that Gln supplementation improves RBC hydration status, a potential mechanistic role that warrants more investigation specifically in sickle RBCs. It is long appreciated that RBCs respond to exogenous metabolites and that the lack of necessary exogenous metabolites negatively impacts RBC lifespan^45, 46^. However, prior to this study, it was unknown how pre-exposure to exogenous amino acids impacted RBCs in the context of oxidative stress and malaria bystander effect. Our study design of amino acid supplementation had an overall focus of recapitulating the plasma environment if key metabolites were supplemented in advance of infection or oxidative stress. Indeed, we found that RBCs supplemented with QCG amino acids are equipped to counter oxidative stress from both H_2_O_2_ and *Pf*CM. We showed that this benefit was intrinsic to RBCs and that ROS development was mitigated within minutes in the cell. As intracellular GSH synthesis occurs in the order of hours within RBCs^47, 48^, this suggests that GSH stores increase in response to QCG pre-incubation, and are not a combative cellular response to oxidative stress. Collectively, our results highlight a beneficial role for exogenous QCG to RBCs prior to oxidative stress. These findings may suggest a prophylactic or therapeutic role of amino acid supplementation in oxidative anemias, including in malaria.

## Acknowledgments

This research was supported in part by federal funds from the National Institute of Health grants 1K01HL143112 through National Heart, Lung and Blood Institute (RJC) and T32 GM127261 (HCB) through National Institute of General Medical Sciences. This research was also supported from awards from Wake Forest University Center for Molecular Signaling and the Wake Forest University School of Medicine Center for Redox Biology and Medicine. The authors wish to acknowledge the support of the Wake Forest Baptist Comprehensive Cancer Center Proteomics and Metabolomics Shared Resource, supported by the National Cancer Institute’s Cancer Center Support Grant award number P30 CA012197. The Microscopic Imaging Core of the Wake Forest Department of Biology provided imaging support for this study. Graphical images were created with BioRender.com.

---

**Supplemental Figure 1:**
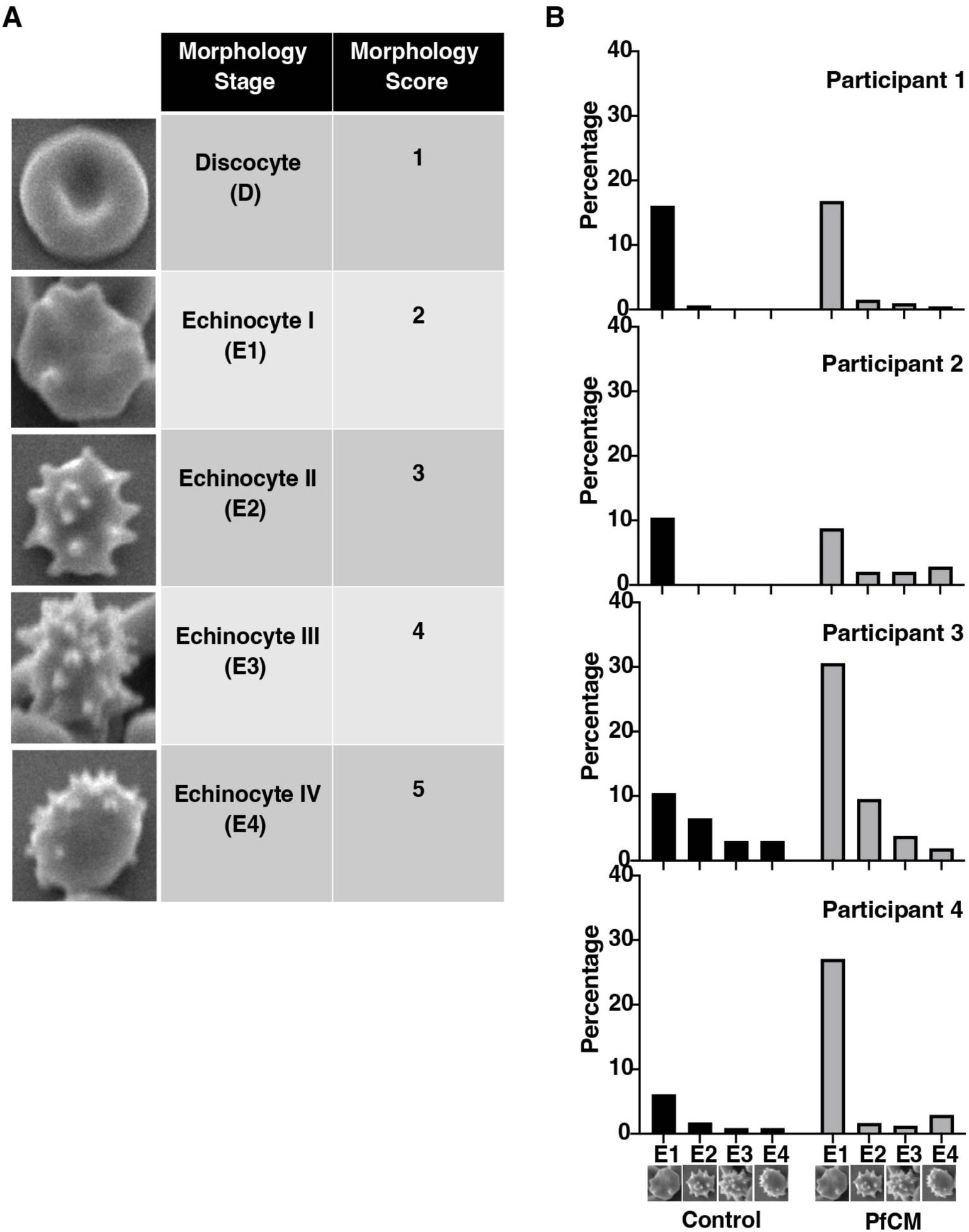
RBC morphology scores and distribution. (**A**) RBCs were fixed and imaged using scanning electron microscopy (SEM) and given a score based on morphology stage. (**B**) Distribution of echinocyte morphologies in RBCs exposed to either control or spent media; each graph indicates an independent participant, n=4.

**Supplemental Figure 2:**
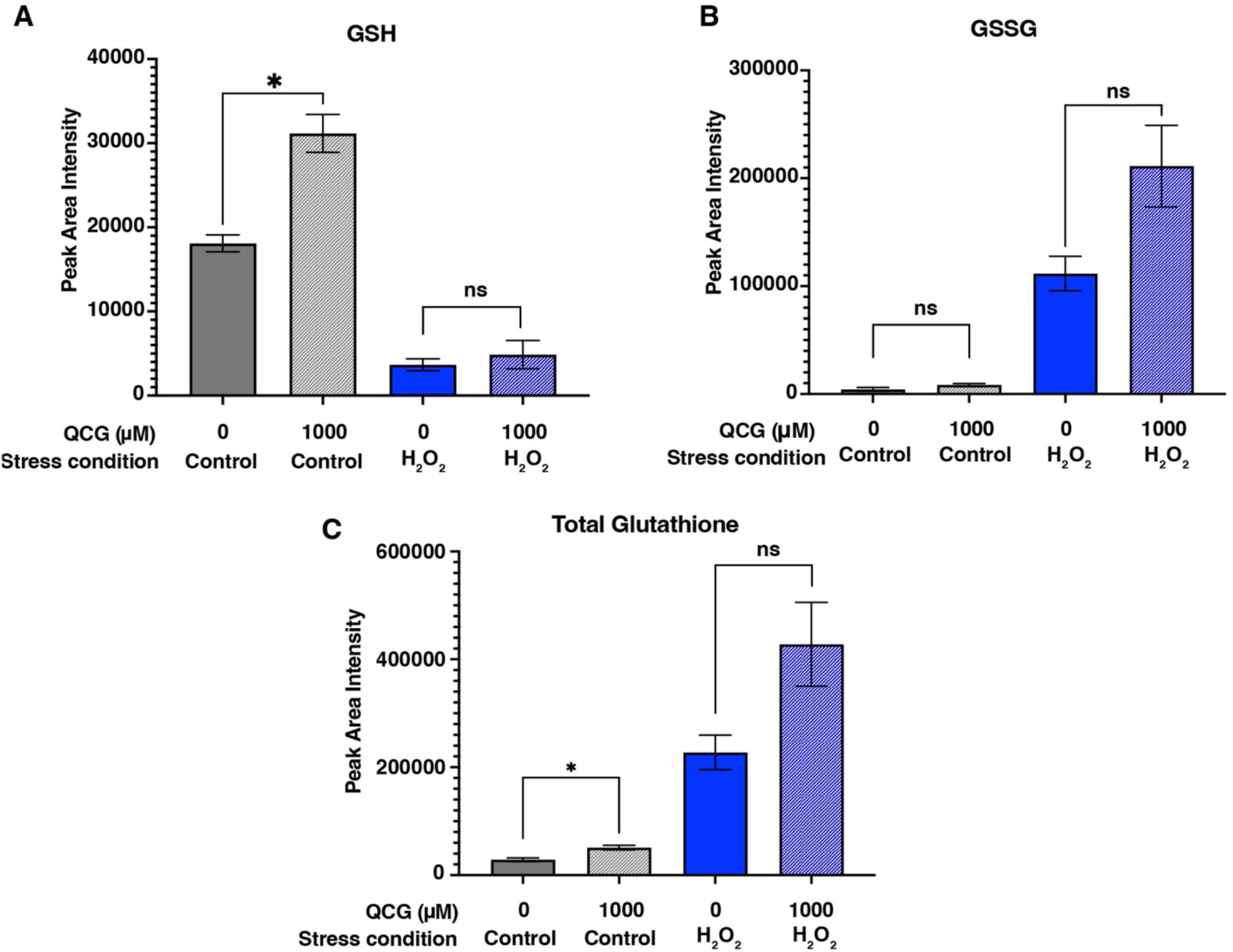
Metabolic changes within RBC as a result of QCG pretreatment followed by H_2_O_2_ stress. Peak area intensity of (**A**) reduced glutathione (GSH), (**B**) oxidized glutathione (GSSG), and (**C**) total glutathione as measured by LC-MS/MS; n=3. One-way paired t-test; ns = not significant, \**p < 0.05*

